# Ethnically relevant consensus Korean reference genome towards personal reference genomes

**DOI:** 10.1101/070805

**Authors:** Yun Sung Cho, Hyunho Kim, Hak-Min Kim, Sungwoong Jho, JeHoon Jun, Yong Joo Lee, Kyun Shik Chae, Chang Geun Kim, Sangsoo Kim, Anders Eriksson, Jeremy S. Edwards, Semin Lee, Byung Chul Kim, Andrea Manica, George M. Church, Jong Bhak

**Author notes:** These authors contributed equally to this work (Yun Sung Cho, Hyunho Kim). These authors jointly supervised this work (George M. Church, Jong Bhak). Correspondence and requests for materials should be addressed to J.B. or to G.M.C.

## Abstract

Human genomes are routinely compared against a universal reference. However, this strategy could miss population-specific or personal genomic variations, which may be detected more efficiently using an ethnically-relevant and/or a personal reference. Here we report a hybrid assembly of Korean reference (KOREF) as a pilot case for constructing personal and ethnic references by combining sequencing and mapping methods. KOREF is also the first consensus variome reference, providing information on millions of variants from additional ethnically homogeneous personal genomes. We found that this ethnically-relevant consensus reference was beneficial for efficiently detecting variants. Systematic comparison of KOREF with previously established human assemblies showed the importance of assembly quality, suggesting the necessity of using new technologies to comprehensively map ethnic and personal genomic structure variations. In the era of large-scale population genome projects, the leveraging of ethnicity-specific genome assemblies as well as the human reference genome will accelerate mapping all human genome diversity.

## Introduction

The standard human reference (currently GRCh38), which is mostly based on individuals of Caucasian and African ancestry^1,2^, is accurate, precise, and extensive. Because of the relatively small long term effective population size of anatomically modern humans (estimated to be as small as ~10,000)^3,4^, this reference is adequate for most purposes and routinely used in research and biomedical applications. However, certain population specific variants could be missed with such a universal reference, and the current research efforts to map human diversity, including low frequency and structural variants, would also benefit from ethnically relevant references^5,6^. Since the publications of the first draft of the human reference genome in 2001^7^, sequencing technologies have advanced rapidly, and additional genome assemblies have been published. In 2007, the diploid genome of a Caucasian male was sequenced and assembled by Sanger technology (HuRef)^8^. Later, the genomes of a Chinese (YH), an African (2009), a Caucasian (HsapALLPATHS1, here called NA12878_Allpaths, 2011), and a Mongolian (2014) were built using Illumina short-read sequencing data only^9-11^. In 2014, a complete hydatidiform mole genome (CHM1_1.1) was assembled, albeit reference-guided, using Illumina short-reads and indexed bacterial artificial chromosome (BAC) clones^12^. In 2015, a haplotype-resolved diploid YH genome was assembled using fosmid pooling together with short-read sequence data^13^. These assemblies, although useful and important for genomics researches, are not of sufficient accuracy or overall quality to be considered a general purpose standard reference genome^14^.

The recent increased availability of long-range sequencing and mapping methods has important implications for the generation of references for ethnic groups and even personal genomes, especially for disease associated structural variations (SVs). Long range data can improve draft genome assemblies by increasing the scaffold size, efficiently closing gaps, resolving complex regions, and identifying SVs^15-22^ at relatively low costs. Notable approaches are single-molecule real-time sequencing technology (SMRT), and highly-parallel library preparation and local assembly of short reads (synthetic long reads) for resolving complex DNA regions and filling genomic gaps^15-17^. For instance, a single haplotype human genome was constructed using single-molecule long read sequencing (CHM1_PacBio_r2, not yet published). Long-read methods can be complemented and validated by two high-throughput mapping methods: optical mapping and nanochannel-based genome mapping. The most representative case is the NA12878 genome (ASM101398v1), which was hybrid assembled by combining single-molecule long reads with single-molecule genome maps (here called NA12878_single)^22^. Assemblies incorporating high-throughput short reads and long range mapping or sequencing data, or hybrid assemblies, can enhance the quality, providing much longer scaffolds with validation and adjustment of complex genomic regions^20-22^.

Complementary to reference genome projects, which provide accurate templates, population genome projects, such as Personal Genome Project (PGP)^23,24^ and the 1,000 Genomes Project (1KGP)^25,26^, provide valuable variome information that is fundamental to many biomedical research projects. PGP was initiated in 2005 to publicly share personal genome, health, and trait data, crucial in understanding the diverse functional consequences associated with genetic variation. Recently, large scale population genome projects in Britain and the Netherlands have been launched to identify population-specific rare genetic variations and disease-causing variants^27,28^. The single reference and population derived genomic variation types and frequencies (variome) are the two main foundations of genomics.

Here, we report a high-quality consensus Korean reference genome (KOREF; reference + variome), produced as part of PGP, by utilizing hybrid sequencing and mapping data. KOREF provides another high quality East-Asian reference to complement GRCh38. KOREF was initiated by the Korean Ministry of Science and Technology in 2006 to generate a national genome reference. To deal with the issues inherent to short reads, we used data from a number of different technologies (short and long paired-end sequences, synthetic and single molecule long reads, and optical and nanochannel genome maps) to build a high quality hybrid assembly (Fig. 1). Furthermore, we integrated information from 40 high-coverage whole genomes (based on short reads) from the Korean PGP (KPGP)^29^ to generate a population-wide consensus Korean reference. We compared the genomic structure of KOREF with other human genome assemblies, uncovering many structural differences, including ethnic-specific highly frequent structural variants. Importantly, the identification of SVs was largely affected by the sequencing platform and assembly quality, suggesting the necessity of long-read sequences and a higher quality assembly to comprehensively map the ethnic and personal genomic structures. Accompanied by multi-ethnic PGP data, in the future, many low-cost personal and ethnic genome references will accelerate the completion of mapping all human genome diversity in both single nucleotide variations (SNVs) and SVs.

**Figure 1.**
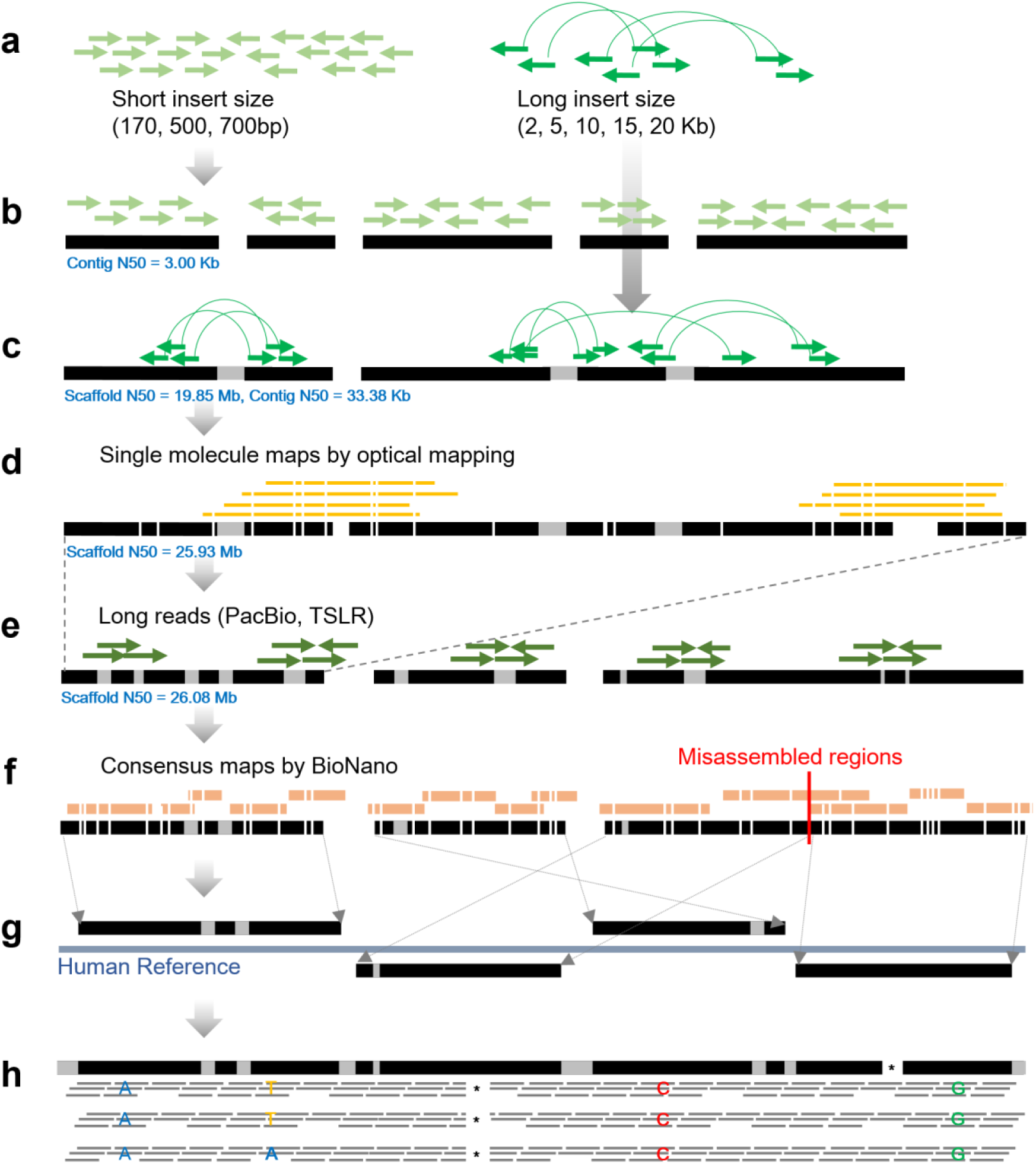
Schematic overview of KOREF assembly procedure. (**a**) Short and long insert size libraries by Illumina whole genome sequencing strategy. (**b**) Contig assembly using *K*-mers from short insert size libraries. (**c**) Scaffold assembly using long insert size libraries. (**d**) Super-scaffold assembly using OpGen whole genome mapping approach. (**e**) Gap closing using PacBio long reads and Illumina TruSeq synthetic long reads (TSLR). (**f**) Assembly assessment using BioNano consensus maps. (**g**) Chromosome sequence building using whole genome alignment information into the human reference (GRCh38). (**h**) Common variants substitution using 40 Korean whole genome sequences.

## Results

### Choosing a representative genome donor

We recruited 16 Korean volunteers, who signed an informed consent (based on the PGP protocol, with minor country-specific adaptations) for use of their genomic data and agreed to their public release. After extracting DNA from peripheral blood (Supplementary Table 1), we genotyped each volunteer using Infinium omni1 quad chip. A multidimensional scaling (MDS) plot of pairwise genetic distances was constructed, using for comparison an additional 34 Korean whole genome sequences from the KPGP database, as well as 86 Japanese, 84 Chinese, 112 Caucasians, and 113 Africans genotype data from HAPMAP phase 3^30^ (Supplementary Fig. 1). All 16 Korean samples fell into a tight population cluster, indicating they are representative of their ethnic group. A healthy male donor was chosen as KOREF by considering a list of parameters such as centrality of the genetic distance, the participant's age, parental sample availability, the availability for continuous blood sample donation, and normality of the G-banded karyotype (Supplementary Fig. 2). In order to supply reference material, an immortalized cell line was constructed from the KOREF donor's blood and deposited in the Korean Cell Line Bank (KCLB, #60211).

### Korean reference genome assembly

We obtained short-read sequencing data from the Illumina HiSeq2000 and HiSeq2500 platforms, using the same approach adopted by other draft reference genome projects^9-11,13,31^. A total 964 Gb of paired-end DNA reads were generated from 24 libraries with different fragment sizes (170bp, 500bp, and 700bp of short insert size, and 2 Kb, 5 Kb, 10 Kb, 15 Kb, and 20 Kb of long insert size), giving a total sequencing depth coverage of ~311 fold (Supplementary Tables 2 and 3). From a *K*-mer analysis, the size of KOREF was estimated to be ~3.03 Gb (Supplementary Table 4). A total of 68,170 scaffolds (≥ 200bp) were generated, totaling 2.92 Gb in length. The assembly reached an N50 length of almost 20 Mb (19.85 Mb) and contained only 1.65% of gaps (Table 1 and Supplementary Fig. 3). Approximately, 90% of the genome draft (N90) was covered by 178 scaffolds, each larger than 3.09 Mb, with the largest scaffold spanning over 80 Mb (81.9) on Chromosome 6.

**Table 1.** KOREF build statistics along the assembly steps

|  | Contig |  | Scaffold |  | Whole-genome<br>optical mapping |  | Long reads<br>(PacBio and TSLR) |  | Chromosomes<br>(Assessment using<br>BioNano maps)<br>*Unplaced scaffolds<br>were excluded. |  |
| --- | --- | --- | --- | --- | --- | --- | --- | --- | --- | --- |
|  | Size<br>(Kb) | No. | Size<br>(Mb) | No. | Size<br>(Mb) | No. | Size<br>(Mb) | No. | Size<br>(Mb) | No. |
| N90 | 8.59 | 89,240 | 3.09 | 178 | 3.86 | 140 | 3.53 | 143 | 81.54 | 19 |
| N80 | 14.62 | 63,987 | 6.45 | 116 | 9.45 | 92 | 9.26 | 93 | 103.05 | 16 |
| N70 | 20.42 | 47,417 | 10.45 | 81 | 14.47 | 67 | 14.53 | 67 | 136.43 | 13 |
| N60 | 26.58 | 35,099 | 16.16 | 59 | 19.56 | 49 | 19.36 | 50 | 137.59 | 11 |
| N50 | 33.38 | 25,446 | 19.85 | 42 | 25.93 | 36 | 26.08 | 36 | 155.88 | 8 |
| Longest | 334.16 | - | 81.91 | - | 101.22 | - | 101.48 | - | 251.92 | - |
| Gaps | 0 % | - | 1.65 % | - | 1.75 % | - | 1.06 % | - | 9.44 % | - |
| Total<br>(≥ 200bp) | 2.87 Gb | 230,514 | 2.92 Gb | 68,170 | 2.92 Gb | 68,103 | 2.94 Gb | 68,451 | 3.12 Gb | 24 |
| Total<br>(≥10 Kb) | 2.52 Gb | 82,254 | 2.88 Gb | 1,243 | 2.88 Gb | 1,176 | 2.90 Gb | 1,369 | 3.12 Gb | 24 |

We then improved the assembly using two methods. We first extend scaffold length by using a high-throughput whole-genome optical mapping instrument, as previously suggested^18^. We extracted high molecular weight DNA and generated 745.5 Gb of single-molecule restriction maps (about two million molecules with a 360 Kb average size) from 67 high density MapCards, resulting in 240-fold optical map coverage (Supplementary Tables 5 and 6). In order to join the scaffolds, the single-molecule optical maps were compared to the assembled scaffolds that were converted into restriction maps by *in silico* restriction enzyme digestion. As a result, a total of 67 scaffolds (>200 Kb) were joined (Supplementary Table 7), resulting in an increased scaffold N50 length of 19.85 Mb to 25.93 Mb (Table 1). Second, we generated two types of long reads for KOREF: PacBio SMRT (~31.1 Gb, ~10-fold coverage; Supplementary Fig. 4 and Supplementary Table 8) and Illumina TruSeq Synthetic Long Reads (TSLR, ~16.3 Gb, ~5.3-fold coverage; Supplementary Fig. 5 and Supplementary Table 9). Both types were used simultaneously, resulting in a decrease in gaps from 1.75% to 1.06% of the expected genome size, and a small increase in final scaffold N50 length from 25.93 Mb to 26.08 Mb (Table 1). To test why the long reads did not improve scaffold lengths, we aligned the two types of long reads onto the KOREF assembly (contigs, scaffolds, and super-scaffolds with optical maps). Much larger portions of the long reads (~8.44%) were aligned to the ends of two different contigs that can be used for scaffolding, but only small portions of the long reads (~0.56%) were aligned to different scaffolds and super-scaffolds (Supplementary Table 10). This result indicates that the continuity information of the long reads were overlapping with those of NGS mate-pair sequences (various insert sizes to ~20 Kb). We suspect that the redundant continuity information between the long reads and the mate pairs, and low sequence depths of the long reads were the main reasons for little increase in the scaffold length.

We then worked to correct any scaffolds misassembles^14,16^. We carefully and systematically assessed the quality of KOREF by generating nanochannel-based genome mapping data (~145 Gb of single-molecule maps > 150 Kb) and assembled the mapping data into 2.8 Gb consensus genome maps having an N50 length of 1.12 Mb (Supplementary Table 11). A total of 93.1% of scaffold regions (≥ 10 Kb) were covered by this consensus map, confirming their continuity (Supplementary Fig. 6). To pinpoint misassembles, we manually checked all the alignment results of the consensus genome map (3,216 cases with align confidence ≥ 20) onto KOREF and GRCh38. Seven misassembled regions were detected in KOREF and were split for correction (Supplementary Fig. 6). Next, we conducted a whole genome alignment of KOREF and GRCh38 to detect possible inter- or intra-chromosomal translocations (indicative of misassembled sequences). A total of 280 of the KOREF scaffolds (≥ 10 Kb) covered 93.5% of GRCh38's chromosomal sequences (non-gaps). We found no large scale inter- or intra-chromosomal translocations. Additionally, as a fine-scale assessment, we aligned the short and long read sequence data to the KOREF scaffolds (self-to-self alignment), and 98.85% of the scaffold sequences (> 2 Kb) were covered by more than 20-fold. Finally, we assigned KOREF’s scaffolds to chromosomes using the whole genome alignment information (chromosomal location and ordering information) of the final scaffolds onto GRCh38 chromosomes, thus obtaining KOREF chromosome sequences (~3.12 Gb of total length; Table 1).

### Consensus variants reference construction and genome annotation

Recently, Dewey *et al*. demonstrated much improved genotype accuracy for disease-associated variant loci using major allele reference sequences^5^, which were built by substituting the ethnicity specific major allele (single base substitutions from the 1KGP) in the low-coverage European, African, and East-Asian reference genomes. We followed the approach for KOREF by substituting sequences with both SNVs and small insertions or deletions (indels) that were commonly found in the 40 Korean PGP high-depth (average 31-fold mapped reads) whole genomes. This removes individual specific biases, and thus better represents common variants in the Korean population as a consensus reference (Supplementary Table 12). About two million variants (1,951,986 SNVs and 219,728 indels), commonly found in the 40 high quality short read Korean genome data, were integrated. Additionally, KOREF’s mitochondrial DNA (mtDNA) was independently sequenced and assembled, resulting in a 16,570bp mitogenome that was similar, in structure, to that of GRCh38. A total of 34 positions of KOREF mtDNA were different from that of GRCh38 (Supplementary Table 13). KOREF’s mtDNA could be assigned to the D4e haplogroup that is common in East-Asians, whereas GRCh38 mtDNA belongs to European haplogroup H.

KOREF GC content and distribution were similar to other human assemblies except African assembly, which has the lowest quality among them (Supplementary Fig. 7). We annotated KOREF for repetitive elements by integrating *de novo* prediction and homology-based alignments. Repetitive elements occupied 1.51 Gb (47.13%) of KOREF (Supplementary Table 14), which is slightly less than found in GRCh38 (1.59 Gb). On the other hand, KOREF contained more repeats than the Mongolian genome (1.36 Gb), which was assembled by next-generation sequencing (NGS) short reads only. We predicted 20,400 protein coding genes for KOREF (Supplementary Table 15 and Methods). By comparing KOREF with other human assemblies (GRCh38, CHM1_1.1, HuRef, African, Mongolian, and YH), a total of 875.8 Kb KOREF sequences (≥100bp of fragments) were defined as novel (Supplementary Table 16 and Methods).

### Korean reference genome compared with other human genomes

We assessed the quality of nine publicly-available human genome assemblies (CHM1_PacBio_r2, CHM1_1.1, NA12878_single, NA12878_Allpaths, HuRef, Mongolian, YH_2.0, African, and KOREF) by comparing assembly statistics, and the recovery rates for GRCh38 genome, segmentally-duplicated regions, and repetitive sequences (Table 2, Supplementary Tables 17–19). The results showed that KOREF was more contiguous (26.46 Mb of N50) than any of the short-read based *de novo* assemblies, but comparable to long-read based assemblies (26.83 Mb of N50 for NA12878_single; 26.90 Mb of N50 for CHM1_PacBio_r2); KOREF was hybrid-assembled by compiling heterogeneous sequencing and mapping technologies, however, a majority of KOREF sequences was derived from NGS short reads. However, KOREF’s contig size is small (47.86 Kb of N50 and 17,749 of L50; Supplementary Table 17) compared to long-read based assemblies due to a low amount of continuity information of short reads. KOREF showed a comparable GRCh38 recovery rate with other long-read assemblies (Supplementary Table 18). KOREF recovered duplicated and repetitive regions more efficiently than other short-read based *de novo* assemblies but less than the two PacBio long-read assemblies (Supplementary Table 19). Especially, the higher depth long-read assembly CHM1_PacBio_r2 recovered the most segmentally-duplicated regions, almost as well as GRCh38, indicating that long read information is important to recover such challenging genomic regions. Also, structural polymorphisms between the two haplotypes in a donor is one of the most significant factors for affecting assembly quality^15,32^, and therefore, it is as expected that CHM1_PacBio_r2, a haploid assembly, showed a much better genome recovery for segmentally-duplicated regions than other assemblies using a diploid source. Additionally, we compared the assembly quality by mapping the re-sequencing data of a single haplotype genome (CHM1) to the human assemblies (Supplementary Fig. 8). Ideally, CHM1 should have no heterozygous variants, if the human assembly recovered the entire genome well. CHM1_PacBio_r2 was the most accurate (having the lowest number of heterozygous variants) in resolving the entire human genome, and KOREF was the most accurate among the short-read based assemblies. Still, these results confirm that short-reads based *de novo* assemblies have a reduced power in fully resolving the entire genome sequences accurately^14^.

**Table 2.** Systematic comparison of assembly quality

| Assembly | Total sequence length (bp) | Scaffold or Contig N50 (Mb) / L50 | Segmental duplication length (bp) | Repeat length (bp) | Detected RefSeq genes (intact only) |
| --- | --- | --- | --- | --- | --- |
| GRCh38<sup>C</sup> | 3,209,286,105 | 67.79 / 16 | 212,777,868 (6.63 %) | 1,564,209,365 (48.74 %) | 20,135 |
| KOREF<sup>S,L,M</sup> | 3,211,075,818 | 26.46 / 35 | 149,353,191 (4.65 %) | 1,452,404,484 (45.23 %) | 17,758 |
| CHM1_PacBio_r2<sup>L</sup> | 2,996,426,293 | 26.90 / 30 | 205,559,250 (6.86 %) | 1,541,211,387 (51.43 %) | 17,657 |
| CHM1_1.1<sup>S,B</sup> | 3,037,866,619 | 50.36 / 20 | 157,426,845 (5.18 %) | 1,417,977,130 (46.68 %) | 18,040 |
| NA12878_single<sup>L,M</sup> | 3,176,574,379 | 26.83 / 37 | 168,652,649 (5.31 %) | 1,545,168,387 (48.64 %) | 6,610 |
| NA12878_Allpaths<sup>S</sup> | 2,786,258,565 | 12.08 / 67 | 90,343,965 (3.24 %) | 1,250,655,296 (44.89 %) | 16,995 |
| HuRef<sup>C</sup> | 2,844,000,504 | 17.66 / 48 | 134,317,812 (4.72 %) | 1,411,487,301 (49.63 %) | 16,968 |
| Mongolian<sup>S</sup> | 2,881,945,563 | 7.63 / 111 | 121,384,034 (4.21 %) | 1,399,420,366 (48.56 %) | 17,189 |
| YH_2.0<sup>S</sup> | 2,911,235,363 | 20.52 / 39 | 127,254,909 (4.37 %) | 1,397,013,571 (47.99 %) | 17,125 |
| African<sup>S</sup> | 2,676,008,911 | 0.062 / 11,689 | 55,830,170 (2.09 %) | 968,988,149 (36.21 %) | 9,167 |

Major sequencing and mapping data used in the assembly are marked by superscript letters: NGS short reads, S; long reads, L; genome maps, M; indexed BAC end sequences, B; chain-terminating Sanger sequences; C.

We also conducted gene content assessments by comparing the number of detected RefSeq^33^ protein-coding genes in each human assembly (Table 2 and Supplementary Table 20). The RefSeq genes were the best recovered in CHM1_1.1 (18,040), which was assembled using that reference as a guide. Among the *de novo* assembled genomes, KOREF contained the largest number (17,758) of intact RefSeq genes, even more than long-read based assemblies (~17,657). Notably, NA12878_single genome, which was hybrid assembled by combining single-molecule long reads with genome maps, had the lowest number (6,610) of intact protein-coding genes, even lower than the low quality African genome (9,167). We confirmed that NA12878_single had many frame-shifts in the coding regions. This can be explained by higher error rates of PacBio single-molecule long reads, which could not be corrected by an error correction step due to its low sequencing depth (46× coverage)^22,34^.

### Structural variation comparison

We investigated SVs, such as large insertions, deletions, and inversions, of eight human assemblies by comparing to GRCh38 (since there were no paired-end read data, HuRef was not used in this analysis). Our assessments showed that the assembly quality is determined mainly by sequencing platform (i.e., sequence read lengths), and therefore, we had to consider that mis-assemblies could generate erroneous SVs. There were two Caucasian samples (CHM1 and NA12878) that were assembled using short-read sequences as well as long reads, and therefore, these assemblies are important in analyzing the association between the assembly quality and SV identification. The CHM1 sample’s ethnicity was confirmed to be Caucasian using ancestry-sensitive DNA makers in autosomes^35^ and mitochondrial DNA sequences (Supplementary Fig. 9). SVs that could be derived from possible misassembles were filtered out by comparing the ratio of aligned single-end reads to paired-end reads (S/P ratio) as previously suggested^36^ (see Methods).

A total of 6,397 insertions (> 50bp), 3,399 deletions (> 50bp), and 42 inversions were found in KOREF compared to GRCh38, for a total of 9,838 SVs. They were slightly fewer than those found in the Mongolian (12,830 SVs) and African (10,772 SVs), but much greater than those found in CHM1 and NA12878 assemblies (~5,179 SVs; Table 3, Supplementary Tables 21 and 22). Notably, YH_2.0 (5,027 SVs) had a similar number of SVs to those found in the Caucasian assemblies, rather than in the other Asian assemblies. The length distribution of the SVs found in the all human assemblies showed a similar pattern (Supplementary Figs. 10 and 11), with a peak at the 200-400bp size range, due to *Alu* element insertions and deletions^15,36^. The fractions of SVs in the repeat regions were higher in the short-read based assemblies (69.6~81.9%) than long-read assemblies (67.7~68.7%; Table 3 and Supplementary Table 23). On the other hand, the fractions of SVs in the segmentally-duplicated regions were much higher in the long-read assemblies (21.4~29.0%) than short-read assemblies (3.9~12.6%; Table 3 and Supplementary Table 24).

**Table 3.**
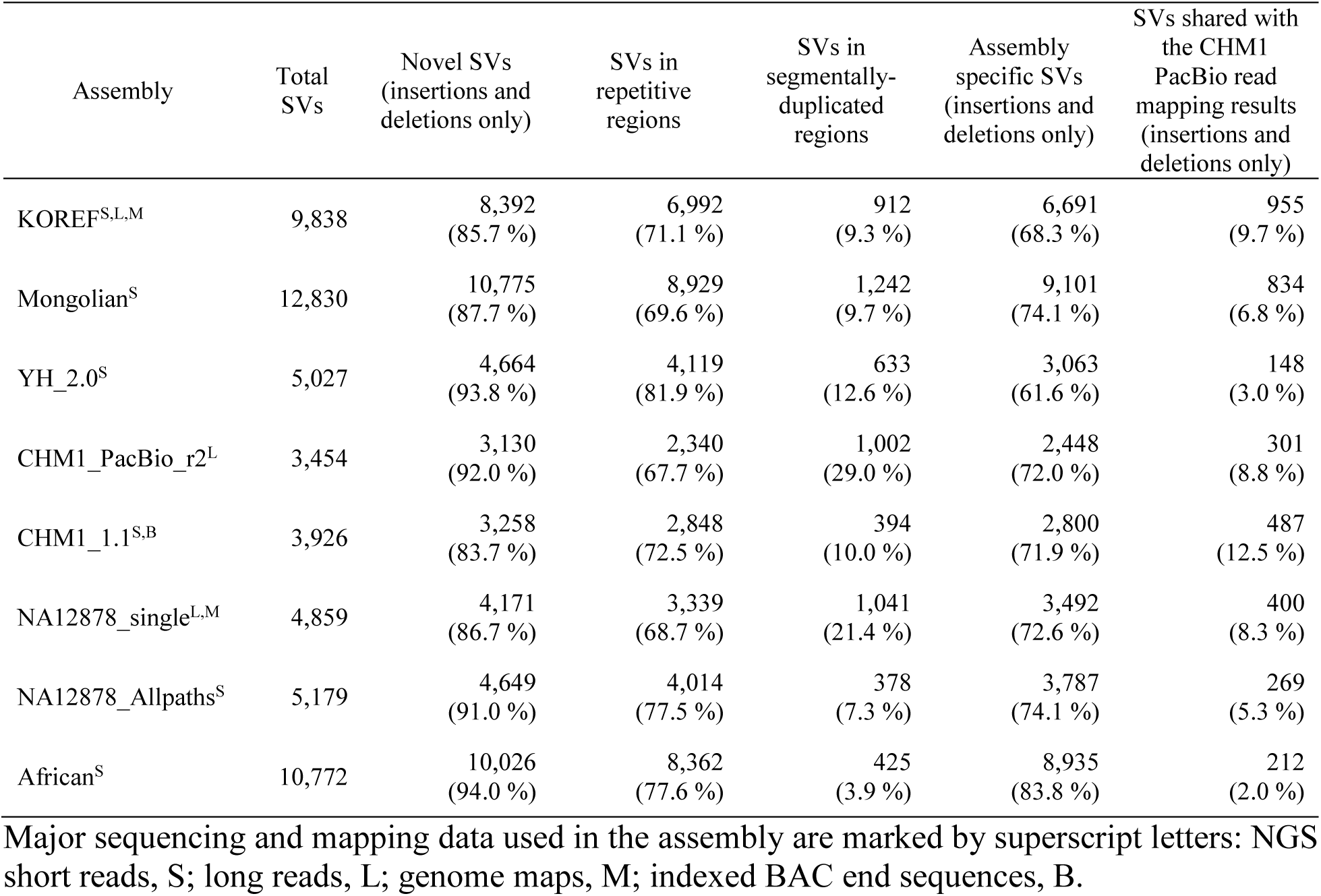
Summary of structural variations in eight human assemblies compared to GRCh38

In KOREF SVs, 93.8% of insertions and 70.4% of deletions were not found in public SV databases and hence defined as novel (Table 3, Supplementary Fig. 10, Supplementary Table 25 and Methods). The fraction of novel SVs in KOREF was similar to those found in other human assemblies but smaller than other short-read only *de novo* assemblies. Regardless of their main sequencing platform, all the assemblies showed a much greater fractions of novel SVs than those found by mapping CHM1’s PacBio SMRT reads to the human reference genome (here called CHM1_mapping)^15^. Notably, CHM1_PacBio_r2, which was assembled using the same sample's PacBio long reads, also showed a much higher fraction of novel SVs. We found a correlation between N50 length of fragments and the fraction of novel SVs (R^2^ = 0.44; Fig. 2a). When we compared SVs of the human assemblies with the SVs by the CHM1_mapping, only small portions of SVs (~12.51%) were shared (Table 3 and Supplementary Table 26). The shared portion of SVs (8.85%) between the CHM1_PacBio_r2 and CHM1_mapping was small, and the shared portions of NA12878 assemblies were quite different (NA12878_single: 8.32%, NA12878_Allpaths: 5.27%). There was a correlation between the assembly quality (N50 length) and shared portion (R^2^ = 0.71; Fig. 2b). These results suggest that even for the same sample there was a large difference between the long-read mapping and *de novo* assembly-based whole genome alignment methods.

**Figure 2.**
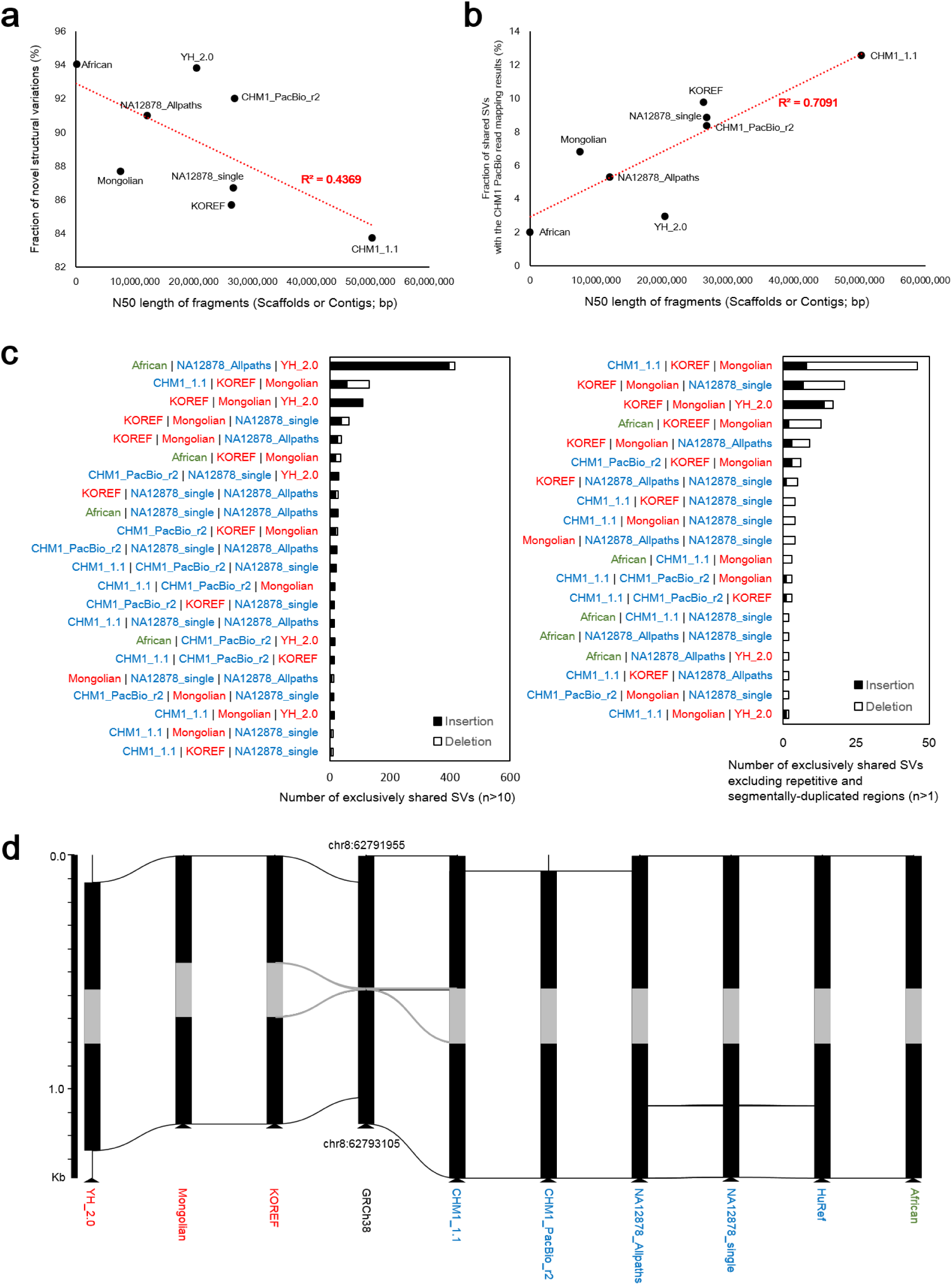
Structural variations among human assemblies. (**a**) The correlation between N50 length of fragments (scaffolds or contigs) and fraction of novel structural variations. (**b**) The correlation between N50 length of fragments and fraction of structural variations shared with the CHM1 PacBio read mapping method. (**c**) Exclusively shared structural variations among human assembly sets. Structural variations shared (reciprocally 50% covered) by only denoted assemblies were considered in this figure. (**d**) An example of structural variation that was shared by nine human assemblies. Gray regions denote structural differences shared among all the assemblies, and horizontal lines indicate homologous sequence regions.

Human genomes contain population-specific sequences and population stratified copy number variable regions^6,37^. Therefore, we assumed that ethnically-relevant human assemblies should share similar genome structures. To investigate the genomic structure among human assemblies, we grouped SVs that were shared by the human assemblies (Fig. 2c). First, most SVs (above 61.6%) were assembly specific (Supplementary Table 27). When we consider SVs that were shared by only two assemblies, two Asian genomes (KOREF and Mongolian) shared the highest number of SVs (Supplementary Fig. 12). However, YH_2.0 shared only small numbers of SVs with KOREF and Mongolian assemblies. Notably, YH_2.0 and African genomes shared SVs abundantly, which cannot be explained by our assumption that similar ethnic genomes should have a higher genome structure similarity. CHM1_PacBio_r2 and NA12878_single, which are Caucasian assemblies using PacBio long read sequences, shared more SVs than those between the same sample’s assemblies (NA12878 assemblies and CHM1 assemblies). In cases of SVs that were shared by only three assemblies, African, NA12878_Allpaths, and YH_2.0 had the largest number of shared SVs, whereas the three Asian genomes had a much smaller number of shared SVs (Fig. 2c and Supplementary Fig. 12). However, when SVs detected in the repetitive and segmentally-duplicated regions were excluded, the three Asian assemblies had the largest number of shared insertions, whereas African, NA12878_Allpaths, and YH_2.0 shared no insertions at all (Supplementary Fig. 13). These results indicate that the SV identification was critically affected by the sequencing platform and assembly quality, and we suggest that long-read sequencing methods are necessary to improve the assembly quality and SV identification for the better characterization of genome structural differences.

Pertaining these limitations, we continued to identify commonly-shared SVs by ethnic groups. To do this, we checked S/P ratios for the SVs using the whole genome re-sequencing data from five Koreans, four East-Asians, four Caucasians, and one African, from KPGP, 1KGP, the Human Genome Diversity Project (HGDP)^38^, and the Pan-Asian Population Genomics Initiative (PAPGI). First, we found one SV that was shared by all human assemblies (Fig. 2d). This SV was also commonly found in the re-sequencing data (13 out of the 14 re-sequencing data). Out of the 110 SVs that were shared by the three Asian assemblies, 18 were frequently found in eleven Asian genomes (one Mongolian assembly, one Chinese assembly, and nine Asian re-sequencing data) compared to ten non-Asian genomes (five non-Asian assemblies and five re-sequencing data, *P*-value <0.05, *Fisher*’s exact test; Supplementary Table 28). Although the SV analysis had limitations due to the heterogeneity of sequencing platform and assembly quality, these results may indicate that the genomic structure is more similar within the same ethnic group^6,37^, suggesting that ethnically-relevant reference genomes are necessary for efficiently performing large-scale comparative genomics.

### Variants comparison mapped to Korean reference genome

Ethnicity-specific genomic sequences that are absent from the reference genome may be important for precise detection of genomic variations^39^. It is also known that the current human reference sequence contains both common and rare disease risk variants^40^, and the use of the current human reference for variant identification may complicate the detection of rare disease risk alleles^5^. Using re-sequencing data on five whole genomes from each population (Caucasian, African, East-Asian, and Korean), we compared the number of variants (SNVs and small indels) detected using KOREF (KOREF single, assembly using single individual; KOREF consensus, assembly after variants substitution by the 40 KPGP genomes) and GRCh38 (Supplementary Tables 29 and 30). We found that the number of variants was significantly different (*P* = 1.04×10^−9^, paired *t*-test), depending on what reference was used (Supplementary Fig. 14). The variant numbers of all individuals (Caucasian, African, and East-Asian) decreased when KOREF consensus was used as a reference. However, because the lower number of actual bases (non-gap) in KOREF could affect the accuracy of genotype reconstruction, we compared variant numbers only within the regions shared by both KOREF and GRCh38 (Supplementary Table 31). As expected, the numbers of homozygous variants from all the Asian genomes (two Chinese, two Japanese, one Mongolian, and five Korean) decreased largely (35.5% of SNVs and 43.9% of indels remained) when KOREF consensus was used as a reference (Fig. 3a and 3b); on the contrary, the numbers of homozygous variants from Caucasian and African genomes decreased little. The numbers of homozygous variants found in non-Korean Asians were similar to those found among Koreans, suggesting that KOREF can be used for other East-Asian genomes. On the other hand, the numbers of heterozygous SNVs were slightly higher in KOREF, which is consistent with the mapping result of the CHM1 re-sequencing data as described above (Supplementary Fig. 8). However, we confirmed that the numbers of heterozygous SNVs became similar to each other when restricting our analysis to non-repetitive regions (data not shown). The numbers of heterozygous indels were also largely constant regardless of the references used (Fig. 3c and 3d).

**Figure 3.**
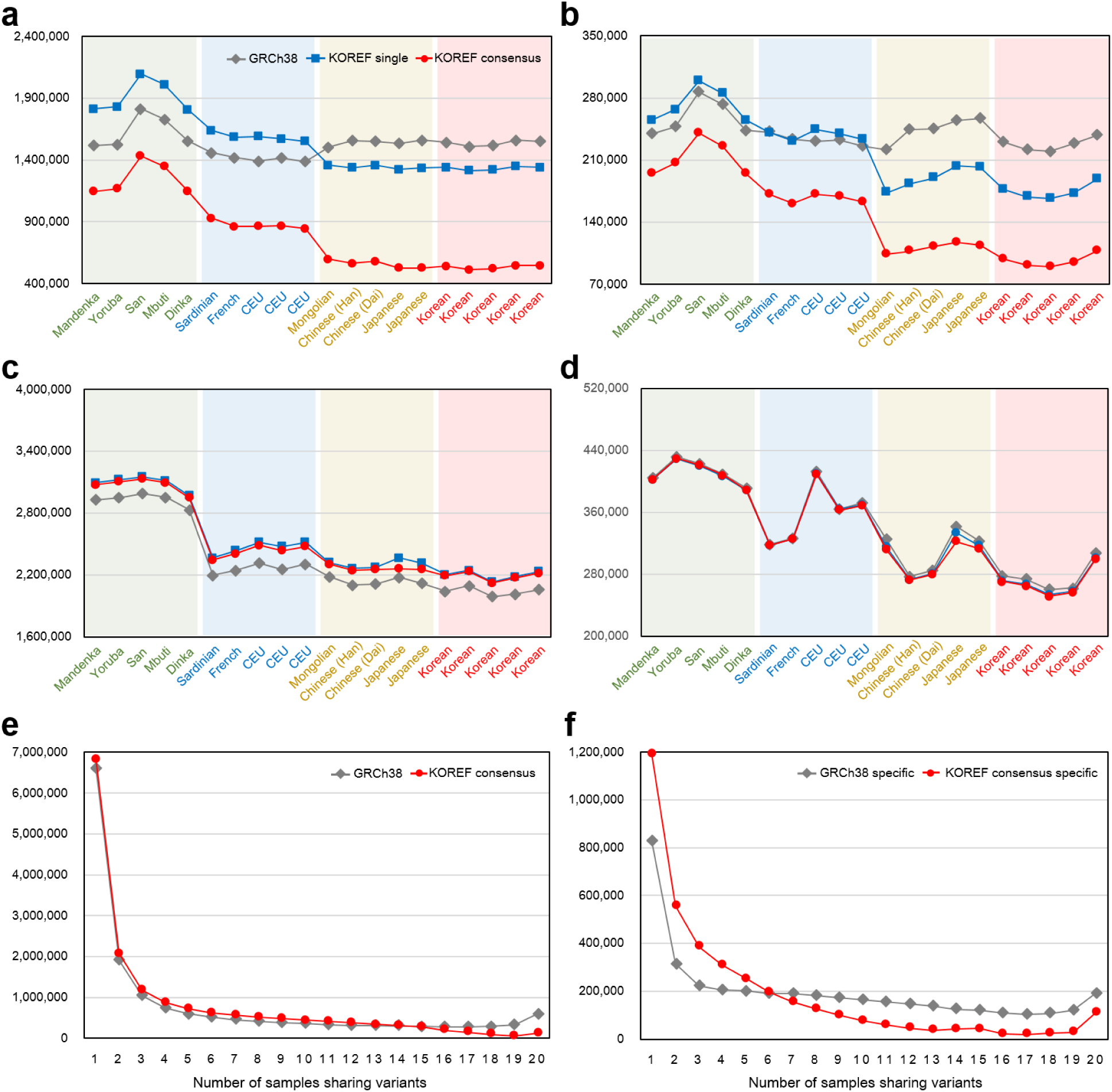
Variants difference depending on the reference genome. Variants (SNVs and small indels) numbers within the regions shared by KOREF and GRCh38 were compared using whole genome re-sequencing data from three different ethnic groups (Africans: Mandenka, Yoruba, San, Mbuti, and Dinka; Caucasians: Sardinian, French, and three CEPH/Utah (CEU); East-Asians: Mongolian, two Chinese, two Japanese, and five Koreans). (**a**) Number of homozygous SNVs. (**b**) Number of homozygous small indels. (**c**) Number of heterozygous SNVs. (**d**) Number of heterozygous small indels. (**e**) The number of variants (referenced by GRCh38 and KOREF consensus) at different levels of sharedness. (**f**) The number of reference-specific variants at different levels of sharedness.

Focusing on differently called variants (variants found in GRCh38 but not found in KOREF consensus, and vice versa), we found that there were differences in the number of variants among populations (i.e., population stratification in terms of variant number). The differences of variants among populations were more prominent when using KOREF specifically called variants (Supplementary Table 32). The number of commonly shared KOREF called variants (> 6 individuals) in the 20 whole genomes was much smaller. Whereas the number of less common KOREF called variants, including individual-specific, was higher (Fig. 3e and 3f). Also, the number of KOREF specifically called variants was significantly lower in the ten Asians than those in the ten non-Asians (*P* = 3.19×10^−10^, *t*-test). These results reflect the consensus variants components of KOREF and also confirm that GRCh38 is depleted for Asian specific sequences^5^. The majority (92.3%) of the GRCh38 specifically called variants were found in dbSNP^41^ (Supplementary Table 32), whereas a smaller fraction (56.17%) of the KOREF specifically called variants were defined as known. When variants in repetitive and segmentally-duplicated regions were excluded, a much larger fraction (86.21%) of the KOREF specifically called variants were known (Supplementary Table 33), indicating that the majority of novel variants found in KOREF was caused by the incompleteness of repetitive and segmentally-duplicated regions. Therefore, we conclude that although KOREF has an advantage for efficient variant detection for the same ethnic genomes especially with the consensus variants components, KOREF needs to be improved using longer sequence reads to reconstruct genotypes properly.

Additionally, we found that the number of variants identified following substitution in the reference with the dominant variant (KOREF single vs. KOREF consensus) is much higher than the change caused by the ethnicity difference (KOREF single and GRCh38; Fig. 3a and 3b). Also, the East-Asians’ homozygous variant number decreased only slightly when the KOREF single was used, compared to GRCh38 (87.0% of homozygous SNVs and 77.3% of homozygous indels remained), while it was greatly decreased when KOREF consensus was used (40.9% of SNVs and 56.8% of indels remained). On the other hand, the number of non-East Asians’ homozygous variants increased when the KOREF single was used, compared to when GRCh38 was used. These results indicate that, at the whole genome variation level, intra-population variation is higher than the inter-population variation in terms of number of variants, supporting the notion that *Homo sapiens* is one population with no genomically significant subspecies.

### Ethnicity-specific reference produces different functional markers

We also found that depending on the reference used, different numbers of non-synonymous SNVs (nsSNVs) and small indels were found in genic regions (Supplementary Tables 34 and 35). With the aforementioned ten East-Asian whole genomes, the number of homozygous nsSNVs (from 3,644 to 1,280 on average) and indels (from 95 to 40 on average) decreased most; whereas a smaller decrease was observed in the five Caucasians (nsSNVs from 3,467 to 2,098; indels from 89 to 65) and five Africans (nsSNVs from 4,216 to 3,007; indels from 134 to 109). When KOREF was used as the reference, predicted functionally altered (or damaged) genes by the homozygous variants also decreased the most among the East-Asians (East Asians, from 490 to 246 on average; Caucasians, from 448 to 362; Africans, from 448 to 415; Supplementary Table 36). Notably, in the ten East-Asians, the functionally altered genes, which were found only against GRCh38 but not KOREF, were enriched in several disease terms (myocardial infarction, hypertension, and genetic predisposition to disease), and olfactory and taste transduction pathways (Supplementary Tables 37 and 38). Additionally, 13 nsSNVs, which are known as disease- and phenotype-associated variants, were called against GRCh38 but not KOREF (Supplementary Table 39); we verified these loci by manually checking short reads alignment to both GRCh38 and KOREF (Supplementary Fig. 15).

## Discussion

In the era of large-scale population genome projects, the leveraging of ethnicity-specific reference genomes as well as GRCh38 could bring additional benefits in detecting variants. This is because each ethnic group has a specific variation repertoire, including single nucleotide polymorphisms and larger structural deviations^6,42^. Population stratification (systematic difference in allele frequencies) can be a problem for association studies, where the association could be found due to the underlying structure of the population and not a disease associated locus^43^. In this study, we provide evidence that an ethnically-relevant consensus reference may improve variant detection. Ethnicity-specific genomic regions such as novel sequences and copy number variable regions can affect precise genotype reconstruction. We demonstrate an example of a better genotype reconstruction KOREF in the copy number variable regions using KOREF (Supplementary Fig. 16). Hence, our ethnicity-specific reference genome, KOREF, may also be useful for detecting disease-relevant variants in East-Asians.

*De novo* assembly based on Sanger sequencing is still too expensive to be used routinely. We have demonstrated that it is possible to produce a *de novo* assembly of relatively high quality at a fraction of the cost by combining the latest sequencing and bioinformatics methods. Additionally, we have shown that optical and nano technologies can extend the size of the large scaffolds while validating the initial assembly. We found that the identification of structural differences based on the genome assembly is largely affected by assembly quality, suggesting a need for new technologies and higher quality of assembly from additional individuals in various populations to better understand comprehensive maps of genomic structure. Also, it is important that the same coordinate system on the GRCh38 allows comparison of different individuals, to leverage the vast amount of previously established knowledge and annotations. Therefore, it is also crucial to research how to transfer those annotations to personal/ethnic reference genomes by preferentially supplementing additional references into GRCh38 in gaining additional biological insights. KOREF cannot, and is not meant to, replace the human reference, and some of its genomic regions, such as centromeric and telomeric regions, and many gaps, are largely incomplete. However, KOREF still can be useful in improving the alignment of East-Asian personal genomes, in terms of fast and efficient variant-calling and detecting individual- and ethnic-specific variations for large-scale genome projects.

## Methods

### Sample preparation

All sample donors in this study have signed the written informed consent. The study has been approved by the Institutional Review Board on Genome Research Foundation (IRB-201307-1 and IRB-201501-1 for KOREF, and 20101202-001 for KPGP). Genomic DNA and RNA used for genotyping, sequencing, and mapping data were extracted from the peripheral blood of sample donors. We conducted genotyping experiments with 16 Korean male participants using Infinium omni1 quad chip to check if the 16 donors had certain genetic biases. A total of 45 Korean whole genomes (40 for variant substitution and five for variant comparison) were used in this study (from the KPGP), sequenced using Illumina HiSeq2000/2500. For the comparison with the 16 donors, 34 Korean whole genome sequences from the KPGP and 86 Japanese, 84 Chinese, 112 Caucasians, and 113 Africans genotyping data from HAPMAP phase 3 were used. After filtering for MAF (< 5%), genotyping rate (< 1%), and LD (R^2^ ≤ 0.2) using PLINK^44^, 90,462 and 72,578 shared nucleotide positions were used to calculate genetic distances for three ethnic groups (East-Asians, Caucasians, and Africans) and three East-Asian groups (Koreans, Chinese, and Japanese), respectively.

Epstein-Barr virus (EBV)-transformed B-cell line was constructed from the KOREF donor’s blood as previously described^45^, with minor modification. Briefly, peripheral blood mononuclear cells (PBMCs) were purified by Ficoll-Paque™ Plus (GE Healthcare, UK) density gradient centrifugation. For EBV infection, the cells were pre-incubated for 1h with spent supernatant from the EBV producer cell line B95-8, and then cultured in RPMI-1640 containing 10-20% fetal bovine serum (FBS), 2mM L-glutamine, 100 U/ml penicillin, 0.1mg/ml streptomycin, 0.25 μg/ml amphotericin B (all from Gibco, Grand Island, NY, USA). The EBV-transformed B-cells were maintained at a concentration between 4 × 10^5^–1 × 10^6^ cells/ml and expanded as needed. The immortalized cell line, named KOREF was deposited in the Korean Cell Line Bank (KCLB, #60211).

### Genome sequencing and scaffold assembly

For the *de novo* assembly of KOREF, 24 DNA libraries (three libraries for each insert size) with multiple insert sizes (170bp, 500bp, 700bp, 2 Kb, 5 Kb, 10 Kb, 15 Kb, and 20 Kb) were constructed according to the protocol of Illumina sample preparation. The libraries were sequenced using HiSeq2500 (three 20 Kb libraries) and HiSeq2000 (others) with a read length of 100bp. PCR duplicated, sequencing and junction adaptor contaminated, and low quality (<Q20) reads were filtered out, leaving only high accurate reads to assemble the Korean genome. Additionally, short insert size and long insert size reads were trimmed into 90bp and 49bp, respectively, to remove poly-A tails and low quality sequences in both ends. Error corrected read pairs by *K*-mer analysis from the short insert size libraries (<1 Kb) were assembled into distinct contigs based on the *K*-mer information using SOAPdenovo2^31^. Then, read pairs from all the libraries were used to concatenate the contigs into scaffolds step by step from short insert size to long insert size libraries using scaff command of SOAPdenovo2 with default options excepting –F option (filling gaps in scaffold). To obtain scaffolds with longest N50 length, we assembled the Korean genome with various *K*-mer values (29, 39, 49, 55, 59, 63, 69, 75, and 79) and finally selected an assembly derived from *K*=55, which has longest contig N50 length. To reduce gaps in the scaffolds, we closed the gaps twice using short insert size reads iteratively.

### Super-scaffold assembly

We used whole-genome optical mapping data to generate a restriction map of the Korean genome and assemble scaffolds into super-scaffolds^18^. First, 13 restriction enzymes were evaluated for compatibility with the Korean genome draft assembly, and *SpeI* enzyme was deemed suitable for the Korean genome analysis. High molecular weight DNA was extracted, and 4,217,937 single molecule restriction maps (62,954 molecules on each map card on overage) were generated from 67 high density MapCards. Among them, 2,071,951 molecules exceeding 250 Kb with ~360 Kb of average size were collected for the genome assembly. The Genome Builder bioinformatics tool of OpGen^18^ was used to compare the optical mapping data to the scaffolds. The distance between restriction enzyme sites in the scaffolds were matched to the lengths of the optical fragments in the optical maps, and matched regions were linked into super-scaffolds. Only scaffolds exceeding 200 Kb were used in this step.

Additionally, we generated two types of long reads for KOREF building: PacBio long reads and TSLRs. The PacBio long reads were generated using a Pacific Biosciences RSII instrument (P4C2 chemistry, 78 SMRT cells; P5C3 chemistry, 51 SMRT cells), and the TSLRs were sequenced by Illumina HiSeq2500. Both long reads were simultaneously used in additional scaffolding and gap closing processes using PBJelly2 program^46^ with default options. To test how much the long reads can contribute to the improvement of scaffolding, the two types of long reads (PacBio and TSLR) were mapped to contigs, scaffolds, and super-scaffolds (using optical maps) of KOREF using BLASR^47^ (version 1.3.1) with default options. To identify reads for scaffolding, we chose best two alignment results by alignment scores. Long reads that were mapped to the ends of two different fragments (allowing a tolerance of 100bp) were considered as reads for scaffolding, if the two alignments shared an overlap below 10% of the read length.

### Assembly assessment and chromosome building

For a large-scale assessment of the scaffolds, we generated nanochannel-based genome mapping data (~145 Gb of single-molecule maps exceeding 150 Kb) on five irysChips and assembled the mapping data into 2.8 Gb of consensus genome maps using BioNano Genomics Irys genome mapping system. The consensus genome maps were compared to KOREF and GRCh38 using irysView software package^22^ (version 2.2.1.8025). To identify misassembles in KOREF in detail, we manually checked alignment results of the consensus genome map into KOREF scaffolds and human reference. For a smaller resolution assessment, we aligned all the filtered short and long reads into the scaffolds using BWA-MEM^48^ (version 0.7.8) with default options. We conducted a whole genome alignment between KOREF scaffolds (≥ 10 Kb) and human reference (soft repeat masked) using SyMap^49^ with default comparison parameters (mapped anchor number ≥ 7) to detect possible inter- or intra-chromosomal rearrangements. We manually checked all the whole genome alignment results.

To build the chromosome sequence of KOREF, first we used the whole genome alignment information (chromosomal location and ordering information) of the final scaffolds (≥ 10 Kb) onto GRCh38 chromosomes. Then, unmapped scaffolds were re-aligned to GRCh38 chromosome with a mapped anchor number ≥ 4 option. Small length scaffolds (from 200bp to 10 Kb) were aligned to GRCh38 chromosomes using BLASR, and only alignments with mapping quality = 254 were used. Unused scaffolds (a total 88.3 Mb sequences) for this chromosome building process were located in an unplaced chromosome (chrUn). Gaps between the aligned scaffolds were estimated based on the length information of the human reference sequences. If some scaffolds’ locations were overlapped, 10 Kb was used as the size of gap between the scaffolds. We added 10 Kb gaps in both sides of KOREF chromosome sequences as telomeric regions as GRCh38 has. The mitochondrial sequences of KOREF were independently sequenced using Nextera XT sample prep kit and then assembled using ABySS^50^ (version 1.5.1) with *K* = 64. Haplogroup of mitochondrial DNA was analyzed using MitoTool^51^.

The 40 Korean whole genome sequences from KPGP database were aligned onto KOREF chromosomes using BWA-MEM with default options, in order to remove individual specific sequence biases of KOREF. SNVs and small indels in the 40 Koreans were called using the Genome Analysis Toolkit (GATK, version 2.3.9)^52^. IndelRealigner was conducted to enhance mapping quality, and base quality scores were recalibrated using the TableRecalibration algorithm of GATK. Commonly found variants in the 40 Korean genomes were used to substitute KOREF sequences. For the SNV substitution, we calculated allele ratio of each position, and then we substituted any KOREF sequence with the most frequent allele only if the KOREF sequence and most frequent allele were different. For the indel substitution, we used only indels that were found in over 40 haploids out of the 40 Korean whole genomes (80 haploids). In cases of sex chromosomes, we used 25 male (25 haploids) whole genomes for Y chromosome and 15 female whole genomes (30 haploids) for X chromosome comparison.

### Genome annotation

KOREF was annotated for repetitive elements and protein coding genes. For the repetitive elements annotation, we searched KOREF for tandem repeats and transposable elements as previously described^10^. For the protein coding gene prediction, homology-based gene prediction was first conducted by searching nucleotides of protein coding genes in Ensembl database 79 against KOREF using Megablast^53^ with identity 95 criterion. The matched sequences were clustered based on their positions in KOREF, and a gene model was predicted using Exonerate software^54^ (version 2.2.0). Also, *de novo* gene prediction was conducted. To certify expression of a predicted gene, we sequenced three different timeline whole transcriptome data of the KOREF sample using a TruSeq RNA sample preparation kit (v2) and HiSeq2500. We predicted protein coding genes with the integrated transcriptome data using AUGUSTUS^55^ (version 3.0.3). We filtered out genes shorter than 50 amino acids and possible pseudogenes having stop-codons. We searched *de novo* predicted genes against primate (human, bonobo, chimpanzee, gorilla, and orangutan) protein sequences from NCBI, and filtered out *de novo* predicted genes if identity and coverage were below 50%. For the assembly quality comparison purpose, we only used homology-based search for RefSeq^33^ human protein-coding genes and repetitive elements. The homology-based segmental duplicated region search was conducted using DupMasker program^56^. To calculate GRCh38 genome recovery rates of human assemblies, we conducted whole genome alignments between each assembly (KOREF final contigs, KOREF final scaffolds, and other assemblies) and GRCh38 using LASTZ^57^ (version 1.03.54) and Kent utilities (written by Jim Kent at UCSC)^58^ with GRCh38 self-alignment options (--step 19 --hspthresh 3000 -- gappedthresh 3000 --seed = 12of19 --minScore 3000 --linearGap medium). After generating a MAF file, we calculated genome recovery rates using mafPairCoverage in mafTools^59^.

To estimate the amount of novel KOREF sequences, we aligned the short insert size and long mate pair library sequences into GRCh38 using BWA-MEM with default options and then extracted unmapped reads using SAMtools^60^ (version 0.1.19) and Picard (version 1.114, http://picard.sourceforge.net) programs. We filtered out possible microbial contamination by searching against Ensembl databases of bacterial genomes and fungal genomes using BLAST with default options. The remaining reads were sequentially aligned into other human genome assemblies (CHM1_1.1, HuRef, African, Mongolian, and YH sequentially) using BWA-MEM with default options, and then removed duplicated reads using MarkDuplicate program in Picard. The alignment results were extracted to an unmapped BAM file using SAMtools view command with -u -f 4 options. We extracted final unmapped reads from the unmapped BAM file using SamToFastq program in Picard. Finally, unmapped reads to the other human genome assemblies were aligned to KOREF. The regions with length ≥100bp and covered by at least three unmapped reads were considered as novel in KOREF.

### Variant and genome comparison

A total of 15 whole genome re-sequencing data results (five Caucasians, five Africans, and five East-Asians) were downloaded from the 1KGP, HGDP, and PAPGI projects. The re-sequencing data (five Caucasians, five Africans, five East-Asians, and five Koreans from KPGP) was filtered (low quality with a Q20 criterion and PCR duplicated reads) and then mapped to KOREF chromosomes with unplaced scaffolds and GRCh38 chromosomes using BWA-MEM with default options. The variants (SNVs and small indels) were called for only chromosome sequences using GATK, in order to exclude variants in unmatched and partially assembled repetitive regions^14^. Variants were annotated using SnpEff^61^, and biological function altering was predicted using PROVEAN^62^. We considered all of the nsSNVs causing stop codon changes and frame shift indels as function altered. Enrichment tests and annotation of variants were conducted using WebGestalt^63^ and ClinVar^64^. The variants were compared with dbSNP^41^ (version 144) to annotate known variants information.

For linking variants found compared to KOREF and GRCh38, the genome to genome alignment was conducted between GRCh38 and KOREF reference genomes using LASTZ^57^. The LASTZ scoring matrix used was with M = 254 (--masking = 254), K = 4500 (--hspthresh = 4500), L = 3000 (--gappedthresh = 3000), Y = 15000 (--ydrop = 15000), H = 0 (--inner = 9), E = 150 / O = 600 (-- gap=<600,150>), and T = 2 options. The LASTZ output was translated to the chain format with axtChain, then merged and sorted by the chainMerge and chainSort programs, respectively. The alignable regions were identified with chainNet, and then selected by netChainSubSet programs for creating a lift-over file. All programs run after LASTZ were written by Jim Kent at UCSC^58^.

To detect SVs among the human genome assemblies, we conducted whole genome alignments between each assembly and GRCh38 using LASTZ. Then, the whole genome alignment results were corrected and re-aligned based on a dynamic-programming algorithm using SOAPsv package. SVs that could be derived from possible misassembles were filtered out by comparing the S/P ratio for each structural variation region in the assembly and GRCh38; authentic SVs would be covered by sufficient paired-end reads, whereas spurious SVs would be covered by wrongly mapped single-end reads. We implemented the S/P ratio filtering system according to the previous published algorithm^36^, because the S/P ratio filtering step in the SOAPsv package is designed for only assembled sequences by SOAPdenovo. *P*-value was calculated by performing Fisher's exact test to test whether the S/P ratio of each SV and the S/P ratio of the whole genome are significantly different (*P*-value < 0.001). We confirmed that commonly shared SVs were not caused by the mis-assembly by checking the mapping status of KOREF short and long reads into both GRCh38 and KOREF. SVs by mapping CHM1's PacBio SMRT reads to the human reference genome were derived by lift-over SV results found against GRCh37 in the published paper^15^. When we compared SVs in the different genome assemblies and available database, we considered SVs to be the same if SVs were reciprocally 50% covered and had the same SV type. Novel SVs were determined as not found in dbVar, Database of Genomic Variants (DGV)^65^, Database of Retrotransposon Insertion Polymorphisms (dbRIP)^66^, dbSNP146, Mills^67^, and 1000 Genome phase 3 database.

## Acknowledgements

This work was supported by the Ministry of Trade, Industry & Energy (MOTIE, Korea) under Industrial Technology Innovation Programs (‘Pilot study of building of Korean Reference Standard Genome map’, No.10046043; ‘Developing Korean Reference Genome’, No.10050164; and ‘National Center for Standard Reference Data’, No.10063239) and Industrial Strategic Technology Development Program (‘Bioinformatics platform development for next generation bioinformation analysis’, No.10040231). This work was also supported by the Korea Research Institute of Bioscience and Biotechnology (KRIBB) under ‘Bioinformatics pipeline construction for de novo assembly’ program. We thank KRIBB people, especially Drs. Tae-Kwang Oh, Woonbong Kim and Kyu-Tae Chang. This work was also supported by ‘Software Convergence Technology Development Program’ through the Ministry of Science, ICT and Future Planning (S0177-16-1046). This work was also supported by the 2015 Research fund (1.150014.01) of Ulsan National Institute of Science & Technology (UNIST). This work was also supported by the Ulsan city. This work was also supported by the Research Fund (14-BR-SS-03) of Civil-Military Technology Cooperation Program. Part of KPGP was supported by KT (Korea Telecom) Personal Genome Project grant. Korea Institute of Science and Technology Information (KISTI) provided us with Korea Research Environment Open NETwork (KREONET) which is the internet connection service for efficient information and data transfer. We thank Mr. Jinup Goh of TheragenEtex for support. We thank INSPUR Co., Ltd., and BIT Co., Ltd. for their technical support. We thank Maryana Bhak for editing.

## Author contributions

J.B. and G.M.C. supervised and coordinated the national Korean reference genome project (KOREF) and Personal Genome Project Korea. J.B., B.C.K., K.S.C., and C.G.K. conceived and designed the reference genome project. J.B. and Y.S.C. coordinated the project's technical research aspects. H.K., H.-M.K., S.J., J.J., and Y.S.C. programmed and performed in-depth data analyses. Cell line construction was performed by Y.J.L. Y.S.C., A.M., G.M.C., and J.B. wrote and revised the manuscript. S.K., A.E., J.S.E., S.L., B.C.K., A.M., and G.M.C. provided critical amendments and edited the manuscript.

## Additional information

### Accession codes

The Korean reference genome project has been deposited at DDBJ/ENA/GenBank under the accession LWKW00000000. The version described in this paper is version LWKW01000000. Raw DNA and RNA sequence reads for KOREF and KPGP have been submitted to the NCBI Sequence Read Archive database (SRA292482, SRA268892).

